# Molluscs community associated with the brown algae of the genus *Cystoseira* in the Gulf of Naples (South Tyrrhenian Sea)

**DOI:** 10.1101/160200

**Authors:** Antonia Chiarore, Sara Fioretti, Angela Meccariello, Giuseppe Saccone, Francesco Paolo Patti

## Abstract

The brown macroalgae of the genus *Cystoseira* are important habitat forming species along the rocky coasts all over the Mediterranean Sea. However, their decline at basin and local scale has been documented in many studies. We have characterized malacofauna associated with *Cystoseira amentacea, C. compressa* and *C. crinita* along the coasts of Ischia Island (Gulf of Naples). Samples were collected by snorkeling in the infralittoral belt. The surface within 20 x 20 cm frames was scraped off and collected in three replicates each sites. The diversity and structure of community were described by number of species, the exponential Shannon and the reciprocal Simpson’s indexes of diversity. The patterns of diversity at spatial scale were assessed by alpha, beta and gamma diversity. A total of 53 species of molluscs were identified in those associations. Gastropoda were the most species-rich class followed by Bivalvia and Polyplacophora. Bivalves were dominant in terms of number of individuals because of the mussel *Mytilus galloprovincialis.* The species *M. galloprovincialis* was the most frequent and top dominant one inhabiting *Cystoseira* associations along the coasts of Ischia Island (96.6 % of the total abundance). Most of the identified molluscs species belonged to two feeding guilds: micrograzers and filter feeders (29 and 13 species respectively). Only juveniles were found providing the importance of *Cystoseira* associations as nursery for molluscs recruitment. Differences in composition and structure of molluscs assemblages were found within the three algal associations and seem to correspond both to different morphology and habitat in which these algal species live.

## Introduction

Marine seaweeds and seagrasses are considered important benthic primary producers along coasts all over the world (Mann 1973). Fucales are the dominant order along the Mediterranean pristine rocky infralittoral shores establishing structurally complex and diversified assemblages and functioning as engineering species (Schiel and Foster 2006). The species of the genus *Cystoseira* together with *Sargassum* are the dominant ones within Fucales in the Mediterranean Sea where most of them are endemic (Giaccone and Bruni 1973) dominating several rocky habitat assemblages from the upper infralittoral shore to the upper circalittoral zone (Ballesteros 1992; Cormaci et al. 2012; Giaccone et al. 1994; Verlaque 1987). They are long-living and very productive macroalgae with a complex tri-dimensional structure providing habitat, food, shelter and nursery for a wide variety of species supporting therefore a high biodiversity (Ballesteros 1992; Bulleri et al. 2002; Mangialajo et al. 2008; Sales et al. 2012; Vergés et al. 2009). They are considered to have an important ecological role also within the European Water Framework Directive (WFD, 2000/60/EC) as coastal water quality indicator (Ballesteros et al. 2007; Orfanidis 2007; Orfanidis et al. 2001). In the last decades, most of *Cystoseira* assemblages in the Mediterranean Sea are suffering a decline or even worse a complete disappearance as an effect of cumulative impacts: habitat destruction, eutrophication, water turbidity, overgrazing by sea urchins, outcompetition by mussels, non-indigenous species and human trampling are considered the major threats (Airoldi et al. 2008; Bianchi et al. 2014; Buia et al. 2013; Cormaci and Furnari 1999; Falace et al. 2010; Giakoumi et al. 2012; Grech et al. 2015; Mangialajo et al. 2008; Sala et al. 2012; Thibaut et al. 2015; Thibaut et al. 2005; Tsiamis et al. 2013). These impacts act over time and in parallel, with a possible synergistic effect on species and ecosystems and on their capability to sustain biodiversity. One of the most evident effect is the replacement of canopy forming algae with less structured and opportunistic species such as turf-forming filamentous seaweeds, mussels or sea urchin barrens involving a simplification of the architectural complexity of the communities (Micheli et al. 2005; Perkol-Finkel and Airoldi 2010; Sala et al. 2012). The loss of habitat structuring species as *Cystoseira* assemblages implies the loss of the associated epibenthic diversity too.

The value of *Cystoseira* associations as a nursery for fish (Cheminée et al. 2013; Lipej et al. 2009; Orlando-Bonaca and Lipej 2005; Riccato et al. 2009; Vergés et al. 2009) as well as the importance in structuring invertebrate communities (Chemello and Milazzo 2002; Fraschetti et al. 2002; Gozler et al. 2010; Milazzo et al. 2000; Pitacco et al. 2014; Urra et al. 2013) have already been investigated in different areas of the Mediterranean Sea.

Amongst the invertebrate fauna inhabiting *Cystoseira* associations mollusca is among the highly represented and dominant taxa, moreover they are considered an important food source for higher trophic levels. In our knowledge, the invertebrate fauna of Gulf of Naples associated with canopy-forming algae of the genus *Cystoseira* has never been investigated. A recent study documented historical changes in this local macroalgal diversity highlighting a drastic decrease of *Cystoseira* species in the infralittoral zones. This decline seems to be associated with the loss of the natural habitat and with the consequent coastal transformation (Grech et al. 2015).

On the other hand, the importance of these algal species in structuring complex and diversified habitat as well as their recent disappearance in the Gulf of Naples, calls for an urgent investigation of these assemblages as well as of the associated fauna biodiversity.

Hence we planned to assess the potential loss of biodiversity associated with these systems in the Gulf of Naples. The characterization of molluscs assemblage structure associated with three *Cystoseira* species along the coasts of Ischia Island, where continuous belt of these algae still persists, was assessed and the pattern of associated diversity at a small spatial scale was studied.

## Materials and methods

### Study sites and sampling design

The study site is Ischia Island, a volcanic island in the south Tyrrhenian Sea. It is located in the northern part of the Gulf of Naples, about 30 kilometres from the city of Naples. It is the largest amongst the Phlegrean Islands. Ischia Island has about 34 km of coastline and a surface area of 46.3 Km^2^. In 2007 was established a marine protected area, Regno di Nettuno, including the Island of Ischia and Procida and the islet of Vivara. The marine protected area Regno di Nettuno is composed by five zones. The zone A in which only relief work and surveillance, service activities and scientific research can be performed on behalf of the managing entity under a specific authorization. In the zone B are possible all the activities allowed in the zone A, bathing, underwater guides tours and diving organized by diving centers, sailing and fishing under specific restrictions. The zone B n.t. (no take) is a zone with particular limitation where professional fishing sports practiced by any means, aquaculture and mussel farming, scuba diving with breathing apparatus are allowed with authorized diving centers. The zone C and D are supervised by rules that allow the recreational use in line with the requirements of eco-compatibility. The morphology of the coast is heterogeneous and it is strictly subject to the geological history of this island. Generally it is possible to define four main geographic sectors identified by the cardinal points. The eastern side is characterized by low rocky coasts and few little sandy beaches. It hosts the biggest harbor of Ischia island that daily connects the island with the mainland. The eastern side of the island falls into the area C of the marine protected area apart from two banks included in the area A.

The morphology of the northern side is similar to that of the eastern part with low rocky coasts and little sandy beaches, this side is characterized by the highest percentage of artificial structure on the coastline, in fact only few scattered individuals of *Cystoseira* species has been detected. The northern coasts fall into the area C of Regno di Nettuno.

The western side is delimited by Punta Caruso and Punta Imperatore. It is characterized by very high rocky coasts and two long sandy beaches. This sector comprises both zones B and C of the marine protected area.

The coast morphology of the southern sector is characterized mainly by high rocky coasts and the largest sandy beach of the island, the Maronti. In this side there is a B n.t zone (the rest of the coasts are under the area B and C.

Six sampling sites along the coasts of Ischia Island were selected according to the previously known presence and co-existence of the assemblages dominated by the three algal species *Cystoseira amentacea, Cystoseira compressa* and *Cystoseira crinita* (Buia et al. 2013). Castello Aragonese – CA, San Pancrazio – SP, Sant’Angelo – SA, Scannella – SC, Punta Imperatore – PI, Punta Caruso – PC. The sampling covered most of the island coastline excepting the northern side where few scattered or even no individuals were recorded. Sant’Angelo falls into the B n.t. zone of the marine protected area, while Scannella and S. Pancrazio into the zone B, the rest of the sampling sites are located in the zone C (Figure 1).

**Figure 1.**
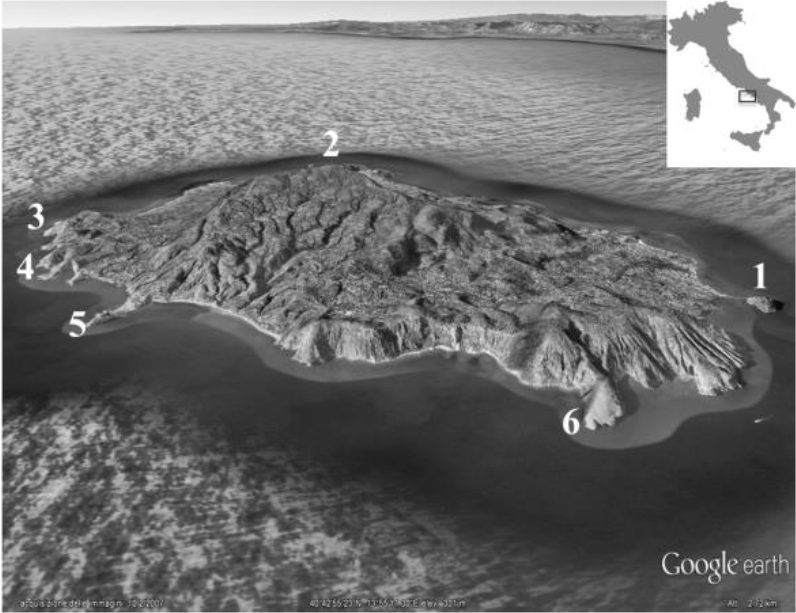
Ischia Island with the six sampling sites. 1: Castello Aragonese – CA; 2: Punta Caruso – PC; 3: Punta Imperatore – PI, 4: Scannella – SC; 5: Sant’Angelo – SA; 6: San Pancrazio – SP.

To avoid the potential bias related to the seasonal variation, the sampling has been carried out during late spring/beginning summer of 2015 and 2016 since this period corresponds with the maximum developmental stage of *Cystoseira* species (Ballesteros 1992; Falace et al. 2004; Hoffmann et al. 1992; Sales and Ballesteros 2012). Moreover, all the chosen sampling sites share some common physical features such as the substrate incline (ranging from 0 to 30 degrees) and the hydrodynamism (mid to high exposed rocky shores). *Cystoseira amentacea* and *Cystoseira compressa* were collected by snorkeling in the upper sublittoral zone (0 meters depth) at the six sampling sites characterized by the co-existence of dense belts of these two algal species. *Cystoseira crinita* was collected by snorkeling only at Scannella (SC) within a tide pool in the proximity of the sea surface since this species was only found at that site. At each site three samples (replicates) were randomly collected by scraping off the macroalgae and associated sessile and vagile fauna within a 20 x 20 cm frame, this area corresponds to the minimum recommended for sampling Mediterranean infralittoral assemblages (Ballesteros 1992; Bianchi et al. 2003a; Boudouresque and Belsher 1979; Coppejans 1980). The number of thalli of macroalgae in each frame was assessed in *situ.* The samples were sealed in individual plastic bag with seawater and preserved in a cool box up to their arrival at the laboratory. The thalli were carefully rinsed in seawater to separate the associated fauna, sorted, the material was sieved through a 0.5 mm mesh and finally the material preserved in 70% absolute ethanol for further taxonomic determination. The maximum height of thalli (the length from the base of the holdfast to the distal tip of the frond) and the dry weight after drying at 60°C for 60 hours were assessed at the laboratory. The molluscs were identified at species level with a stereomicroscope according to Cattaneo-Vietti et al. (1990); Cossignani and Ardovini (2011); Doneddu and Trainito (2005); Giannuzzi-Savelli (2003) and counted. Updated taxonomy and nomenclature was cross-checked with the World Register of Marine Species database WoRMS (Appeltans et al. 2012), last accessed: 30 November 2016. The Check List of European Marine Molluscs CLEMAM (Gofas and Le Renard 2013) was followed for the systematic status of the species, last accessed: 30 November 2016.

According to the feeding guilds as described by Solis-Weiss et al. (2004) and Rueda et al. (2009), the following categories were considered: carnivores feeding on other mobile organisms (C); scavengers feeding on remains of dead organisms (SC); deposit feeders feeding on organic particles contained in the sediment (D); ectoparasites and specialised carnivores feeding on much larger organisms on which they live during their life cycle (E); filter feeders capturing the particles in the water column with their gills and/or with mucous strings (FF); micrograzers feeding on microalgae, cyanobacteria or detritus attached to algal fronds (MG), macroalgae grazers (AG).

### Analysis of macroalgal features

The density as the mean number of thalli per 400 cm^2^, the mean height of thalli in centimeters and the dry biomass after drying for 60hr at 60°C were calculated. Since none of the macroalgal measures (except the mean height) displayed normal distribution (Shapiro-Wilks test) and homoscedasticity (Bartlett test) neither after data transformation, a non-parametric PERMutational multivariate ANalysis Of VAriance, PERMANOVA (Anderson et al. 2008; Anderson 2001a) applied on the Euclidean distance matrix of raw data was chosen to test differences among macroalgal features at the six sites. PERMANOVA design included two factors: alga (fixed factor, 3 levels) and site (random factor, 6 levels). P-values were obtained by 9999 permutations of the raw data under an unrestricted model. Pairwise comparisons for all the combinations of alga x site were also performed using t tests and 9999 permutations of the raw data. In order to avoid the potential lack of analysis robustness to heterogeneity of data for unbalanced design (Anderson and Walsh 2013), a reduced analysis only including data from the site SC where the three algal species occurred simultaneously was also performed, in this case the PERMANOVA design included only the factor alga (fixed, three levels). All the multivariate analyses were carried out by the software PRIMER v 6.1.11 with PERMANOVA + V. 1.0.1 add-on package, developed by the Plymouth Marine Laboratory (Clarke and Gorley 2001). The tests for normality and homoscedasticity of data were performed using R V. 3.2.2 (R-Core-Team 2013).

### Analysis of the associated assemblages structure and species diversity

Data from the two sampling years were analyzed together, as a result at each sites six replicates were considered instead of three. The species were quantified according to: abundance (total number of individuals collected), frequency index (percentage of samples in which a species is present) and dominance index (percentage of individuals of a species within the sample on the total). The diversity patterns and assemblage structure of malacofauna were described through different diversity measures: number of species (S), the exponential Shannon index (ExpH’) and the reciprocal Simpson’s index of diversity (1/Simpson) following the suggestion of Jost et al. (2010) to estimate the ‘effective number of species’. The cumulative ranked species abundance or *k*-dominance curves were performed to extract overall information on pattern of relative species abundance associated with the three algal species. The dominance curves are based on ranking the species in a sample in decreasing order of their abundance, the ranked abundances are expressed as a percentage of the total abundance of all species in the sample, in the case of *k*-dominance curve, the cumulative ranked abundance are used (Clarke 1990). The patterns of diversity at different spatial scales were assessed by analyzing alpha diversity (average number of species per sample unit), gamma diversity (the total number of species within a sampling site) and beta diversity (the changes in species composition between sampling sites). Beta diversity was calculated as the multivariate measure based on the average distance between group-centroids defined by a distance matrix determined with the PERMDISP procedure. PERMDISP is a test used to compare the sample dispersion of different groups based on a distance matrix, when it is applied on a Jaccard distance presence/absence data matrix, it is directly interpretable as a measure of beta diversity among groups (Anderson et al. 2011).

To visualize the spatial pattern of similarity of mollusc assemblages in the three algal species, non-metric multidimensional scaling (nMDS) plot (Kruskal 1964b) was performed on the distance among centroids matrix derived from a Bray-Curtis similarity matrix using square-root-transformed abundance data.

Furthermore a similarity percentage analysis SIMPER (Clarke and Warwick 1994) was performed to identify the species responsible for the similarity/dissimilarity within and between the three algal species at the different sites.

Multivariate approaches were also used to appraise the composition of mollusc assemblages associated with the three algal species. A nonparametric analysis of variance, PERMANOVA (Anderson et al. 2008; Anderson 2001a; Anderson 2001b) applied on a Bray-Curtis similarity matrix using square-root-transformed abundance data in order to down-weight the abundant species was used, the model included two factors: alga (fixed factor, three levels) and sites (random factor, six levels). Pair-wise comparisons for all the combinations of alga x site were also performed using t tests and 9999 permutations of the raw data. PERMANOVA was also performed to test differences in the values of diversity index applied on an Euclidean distance data matrix. All the multivariate analyses were carried out by the software PRIMER v 6.1.11 with PERMANOVA + V 1.0.1 add-on package, developed by the Plymouth Marine Laboratory.

## Results

### Description of macroalgal features

No significant differences were found by comparing data from the reduced analysis; hence, we reported results from the overall analyses. Measures of the macroalgal features are reported in the Table 1 and shown in the Figure 2. No significant density differences were found among sites (F_5,65_ = 0.89, p > 0.05) neither among algae (F_2,65_ = 4.05, p > 0.05). The average height was significantly different both among algae and sites (F_2,65_ = 20.07 p < 0.01 and F_5,65_ = 5.3735 p < 0.001 respectively). Biomass showed significant differences among the six sampling sites (F_5,65_ = 6.42, p < 0.001) with a maximum resemblance distance between *C. compressa* and *C. amentacea* at PI, however no significant differences were found among algae (F_2,65_ = 0.0088, p > 0.05), for further details see Table 2. PERMANOVA results of the pair-wise t-test applied on macroalgal features for the interaction alga x site are reported in the Table 3. *Cystoseira compressa* reaches the highest mean values of density in almost all the sampling site apart from PI and SP where *Cystoseira amentacea* mean values of density are slightly higher. *C. amentacea* reaches the highest mean values of height in all the sampling sites compared to *C. compressa,* however at SC *Cystoseira crinita* has the highest mean value of thalli height. At SP and PI, *C. amentacea* mean value of dry biomass are higher than those of *C. compressa* at the same sampling sites, however at the other sites *C. compressa* reaches higher values of dry biomass. At SC the mean value of biomass are comparable among the three algal species.

**Figure 2.**
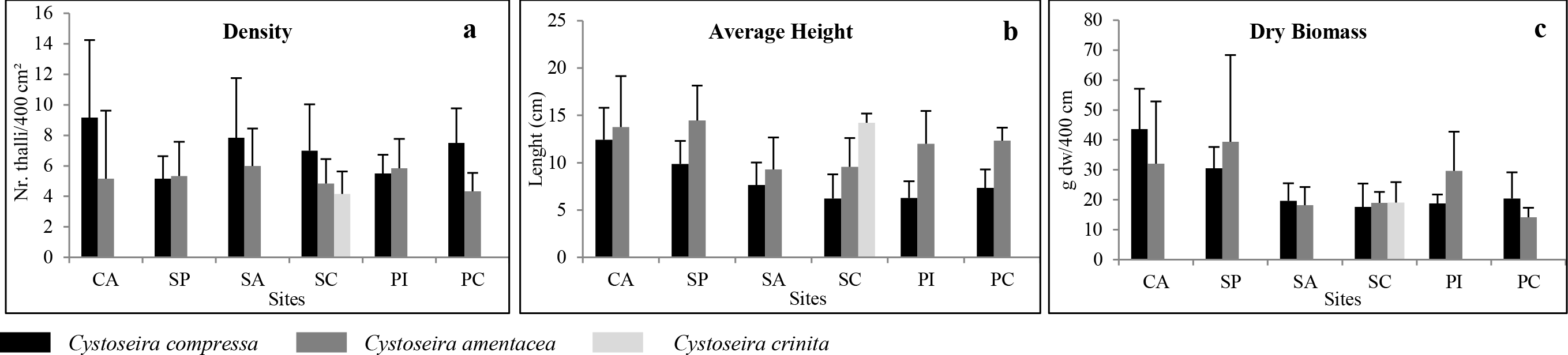
Mean values ± SD of the three macroalgal measures at the six sampling sites. a: density, b: average height, c: dry biomass

**Table 1.** Mean value (± SD) of macroalgal features at the six sampling sites. *Cystoseira compressa: C. co, Cystoseira amentacea: C. am, Cystoseira crinita: C. cr*.

| Site | Density (Nr. Thalli/400 cm<sup>2</sup>) |  |  | Height (cm) |  |  | Biomass (g dw/400 cm<sup>2</sup>) |  |  |
| --- | --- | --- | --- | --- | --- | --- | --- | --- | --- |
|  | *C. co* | *C. am* | *C. cr* | *C. co* | *C. am* | *C. cr* | *C. co* | *C. am* | *C. cr* |
| CA | 9.2 ± 5.1 | 5.2 ± 4.5 | ----- | 12.4 ± 3.4 | 13.8 ± 5.4 | ----- | 43.6 ± 13.5 | 32.0 ± 20.8 | ----- |
| SP | 5.2 ± 1.5 | 5.3 ± 2.2 | ----- | 9.9 ± 2.5 | 14.5 ± 3.7 | ----- | 30.5 ± 7 | 39.3 ± 29.0 | ----- |
| SA | 7.8 ± 3.9 | 6.0 ± 2.4 | ----- | 7.7 ± 2.4 | 9.3 ± 3.4 | ----- | 19.6 ± 5.8 | 18.2 ± 6.1 | ----- |
| SC | 7.0 ± 3.0 | 4.8 ± 1.6 | 4.2 ± 1.5 | 6.2 ± 2.5 | 9.6 ± 3.0 | 14.2 ± 3.6 | 17.7 ± 7.7 | 19.0 ± 3.7 | 19.1 ± 6.8 |
| PI | 5.5 ± 1.2 | 5.8 ± 1.9 | ----- | 6.3 ± 1.7 | 12.0 ± 3.5 | ----- | 18.8 ± 3.0 | 29.7 ± 13.1 | ----- |
| PC | 7.5 ± 2.3 | 4.3 ± 1.2 | ----- | 7.3 ± 1.9 | 12.3 ± 1.4 | ----- | 20.4 ± 8.7 | 14.1 ± 3.2 | ----- |

**Table 2.** Differences among macroalgal features tested with PERMANOVA for the factors alga (fixed, 3 levels) and site (random, 6 levels) and their interaction (alga x site). Pseudo-F values by 9999 permutation. Df: degrees of freedom. Significance: * p < 0.05, ** p < 0.01, *** p < 0.001, ns = not-significant.

| Pseudo F-value |  |  |  |
| --- | --- | --- | --- |
| Variables | Factors |  | Interaction |
|  | Alga (df = 2) | Site (df = 5) | Alga x Site (df = 5) |
| Density | 4.05 ns | 0.89 ns | 1.13 ns |
| Height | 20.01 ** | 5.37 *** | 1.01 ns |
| Biomass | 0.0088 ns | 6.42 *** | 1.48 ns |

**Table 3.** PERMANOVA results of pair-wise t-tests applied on macroalgal features for the interaction alga x site for pair of levels of factor alga. t values by 9999 permutation. Significance: * p < 0.05, ** p < 0.01, *** p < 0.001, ns = not-significant. *a: t test between C. compressa and C. amentacea, b: t test between C. compressa / C. crinita, c: t test between C. amentacea / C. crinita*

| t-values |  |  |  |  |  |  |
| --- | --- | --- | --- | --- | --- | --- |
| Sites | CA | SP | SA | SC | PI | PC |
| **Density** | 1.45 ns | 0.15 ns | 0.97 ns | 1.55<sub>a</sub> ns<br>2.06<sub>b</sub> ns<br>0.75<sub>c</sub> ns | 0.17 ns | 3.03 ** |
| **Biomass** | 1.14 ns | 0.72 ns | 0.42 ns | 0.38<sub>a</sub> ns<br>0.34<sub>b</sub> ns<br>0.01<sub>c</sub> ns | 1.98 * | 1.64 ns |
| **Height** | 0.52 ns | 2.56 * | 0.98 ns | 2.06<sub>a</sub> ns<br>4.43<sub>b</sub> **<br>2.41<sub>c</sub> * | 3.59 ** | 5.13 ** |

### Description of the associated assemblages structure and species diversity

A total of 24837 individuals inhabiting the three associations of *Cystoseira amentacea*, *Cystoseira compressa* and *Cystoseira crinita* along the coasts of Ischia Island were collected. 53 species were identified, belonging to three classes and 31 families, including Polyplacophora (2 families), Gastropoda (19 families) and Bivalvia (10 families). Gastropoda was the class showing the highest number of species (38 species), followed by Bivalvia (13 species) and Polyplacophora (2 species). The best represented families were Rissoidae (10 species) and Phasianellidae (3 species) for gastropods and Mytilidae (3 species) for bivalves. A detailed species list is shown in the Table 4. Bivalvia was the most important class in terms of abundance with 24104 individuals (97 % of the total abundance), followed by Gastropoda: 729 individuals (2.9%) and Polyplacophora: 4 individuals (0.02%). All the individuals were present at juvenile stage. Most of the identified mollusc species belonged to two main feeding guilds: filter feeders (13 species) and micrograzers (29 species). Only 3 species of carnivores were found, 3 species of scavengers, 3 specialized carnivores (three species of nudibranchs), one species of macroalgae grazers (*Aplysia punctata*) and only one species belongs to deposit feeders (*Scissurella costata).*

**Table 4.**
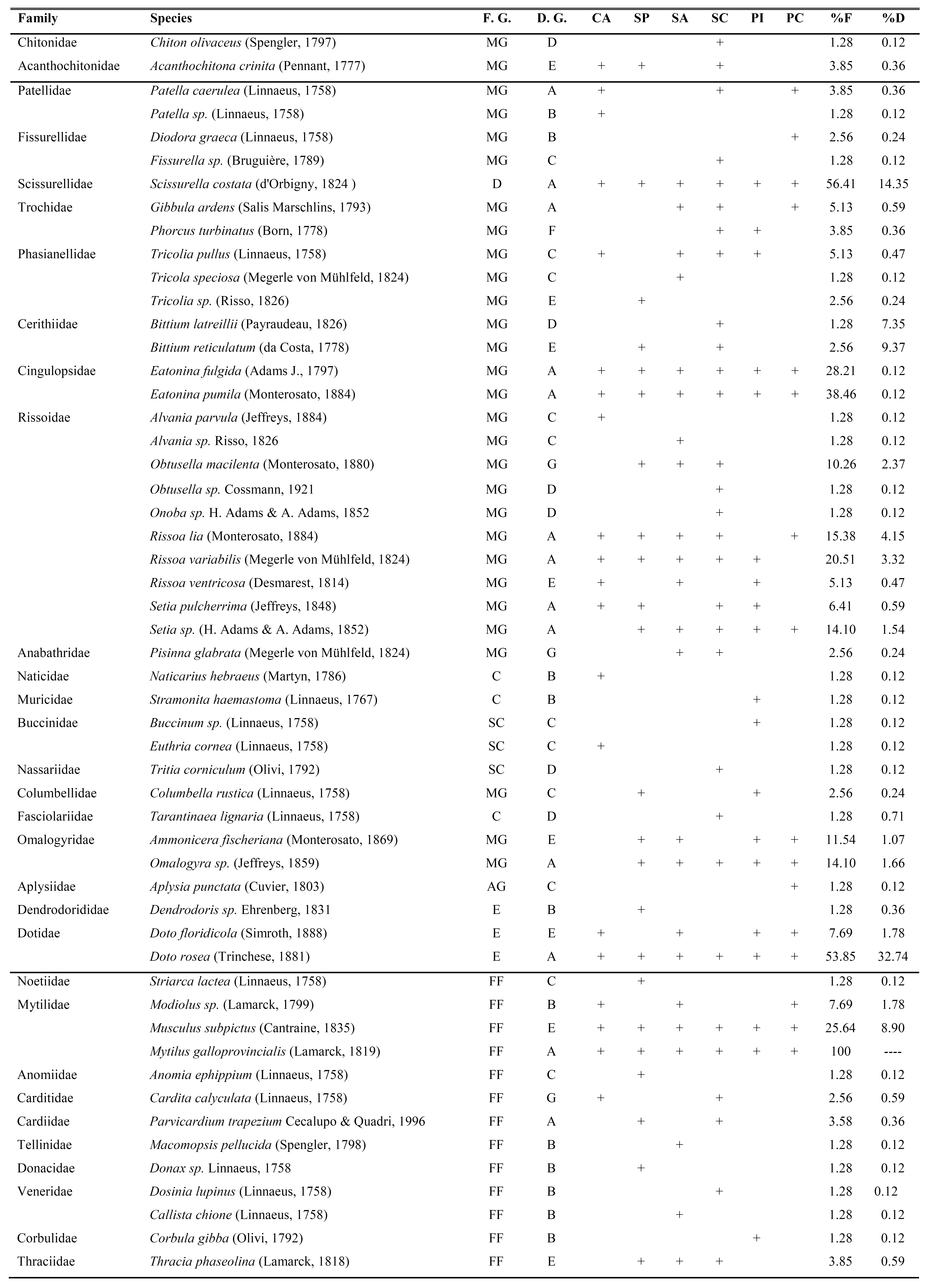
List of the species identified within the three algal associations at the six sampling sites in systematical order. Feeding Guild (F. G.): micrograzers (MG); filter feeders (FF); carnivores (C); deposit feeders (D); scavengers (SC), macroalgae grazers (AG); ectoparasites and specialized carnivore (E). Distribution Group (D. G.): shared species among the three algae (A); exclusive species for *C*. *compressa* (B), exclusive species for *C. amentacea* (C); exclusive species for *C. crinita* (D); shared species *C. compressa/C. amentacea* (E), shared species *C. compressa/C. crinita* (F); shared species *C. amentacea/C. crinita* (G). Sampling sites: Castello Aragonese (CA); San Pancrazio (SP); Sant’Angelo (SA); Scannella (SC); Punta Imperatore (PI), Punta Caruso (PC). +: present; empty: absent;

The species *Mytilus galloprovincialis* was ubiquitously found within the three algal assemblages at the six sampling sites, moreover it was the most important species in terms of abundance with a total of 23994 individuals contributing to the 96.6% of the total abundance of all the individuals. In order to avoid the potential homogenization of mollusc community biodiversity due to *M. galloprovincialis,* the following analysis did not take into account this species. *Scissurella costata, Eatonina fulgida, Eatonina pumila* and *Doto rosea* were also ubiquitously found within the three algal species at the six sampling sites.

No significant differences were found in the total number of species associated with the three algae (F_2,65_ = 3.95, p > 0.05) neither among sites (F_5,65_ = 0.21, p > 0.05) (Table 5). Differences were found in the number of individuals with a maximum resemblance distance between *C. compressa* and *C. amentacea* at SP (pair-wise test t = 2.4, p < 0.05). PERMANOVA results of pair-wise t-tests applied on diversity index for the interaction alga × site for pair of levels of the factor alga are reported in the Table 6.

**Table 5.** Differences among diversity index tested with PERMANOVA for the factors alga (fixed, 3 levels) and site (random, 6 levels) and their interaction (alga × site). Pseudo-F values by 9999 permutation. Df: degrees of freedom. Significance: * p < 0.05, ** p < 0.01, *** p < 0.001, ns = not-significant. Exp H’= exponential Shannon, 1/Simpson= inverse Simpson

| Variables | Pseudo F-values |  |  |
| --- | --- | --- | --- |
|  | Alga (df = 2) | Site (df = 5) | Alga x Site (df = 5) |
| **Nr. Species** | 3.96 ns | 0.21 ns | 1.53 ns |
| **Nr. Individuals** | 2.085 ns | 1.58 ns | 2.85 ** |
| **Exp. H'** | 10.88 * | 0.66 ns | 1.01 ns |
| **1/Simpson** | 11.49 * | 0.73 ns | 0.53 ns |

**Table 6.** PERMANOVA results of pair-wise t-tests applied on macroalgal features for the interaction alga × site for pair of levels of factor alga. t values by 9999 permutation. Significance: * p < 0.05, ** p < 0.01, *** p < 0.001, ns = not-significant. *a: t test between C. compressa and C. amentacea, b: t test between C. compressa / C. crinita, c: t test between C. amentacea / C. crinita.*

| t-values |  |  |  |  |  |  |
| --- | --- | --- | --- | --- | --- | --- |
| Sites | CA | SP | SA | SC | PI | PC |
| **Nr. species** | 0.35 ns | 2.29 ns | 0.86 ns | 1.04<sub>a</sub> ns<br>2.80<sub>b</sub> *<br>1.97<sub>c</sub> ns | 0.33 ns | 1.46 ns |
| **Nr. individuals** | 1.03 ns | 2.36 ns | 1.74 ns | 0.75<sub>a</sub> ns<br>0.91<sub>b</sub> ns<br>0.18<sub>c</sub> ns | 1.22 ns | 0.44 ** |
| **Exp H'** | 0.33 ns | 1.20 ns | 0.38 ns | 0.75<sub>a</sub> ns<br>2.50<sub>b</sub> *<br>2.06<sub>c</sub> ns | 0.40 ns | 1.98 ns |
| **1/Simpson** | 0.46 ns | 0.60 ns | 0.24 ns | 0.18<sub>a</sub> ns<br>1.99<sub>b</sub> **<br>1.92<sub>c</sub> * | 0.82 ns | 2.18 ns |

The species associated with *C. compressa* were 33 (22 Gastropoda, 10 Bivalvia and 1 Polyplacophora), representing 24.4% of the total abundances. The same number of species was associated with *C. amentacea* (27 Gastropoda, 5 Bivalvia and 1 Polyplacophora) with a contribution of 67% to the total number of individuals. The number of species associated with *C. crinita* were 22 (19 Gastropoda, 2 Bivalvia and 1 Polyplacophora) with a contribution of 8.6 % to the total abundances.

The three algal assemblages shared a total of 12 species (11 Gastropoda and 1 Bivalvia). The species exclusively associated with one or two algal species were as follow: 11 species exclusively associated with *C. compressa* (5 Gastropoda and 6 Bivalvia), 11 species only associated with *C. amentacea* (9 Gastropoda, 2 Bivalvia), six species only associated with *C. crinita* (1 Polyplacophora, 5 Gastropoda). Furthemore 8 mollusc species were shared between *C. compressa/C.amentacea* (1 Polyplacophora, 5 Gastropoda, 2 Bivalvia), the species *Phorcus turbinatus* was shared among *C. compressa* and *C. crinita,* finally the species *Obtusella macilenta*, *Pisinna glabrata* and *Cardita calyculata* were shared by *C. amentacea* and *C. crinita*.

The most frequent (58.3%) and abundant (30.6%) species associated with *C. compressa* was *Scissurella costata,* followed by *Eatonina pumila* (39% of frequency, 10.7% of abundance) and *Doto rosea* (30.6% of frequency, 9.7% of abundance). *Doto rosea* was the most frequent (80.6%) and abundant (45%) species associated with *C. amentacea*, followed by *Scissurella costata* (50% of frequency, 8.2% of abundance), *Eatonina pumila* (41.7% of frequency, 9.8% of abundance) and *Musculus subpictus* (39% of frequency, 11.4% of abundance). *Scissurella costata* was also the most frequent species associated with *C. crinita* (83.3%), but the most abundant one was *Obtusella macilenta* (17.8%). Values of alpha, beta and gamma diversity as well as the index of diversity are graphed in the Figure 3.

**Figure 3.**
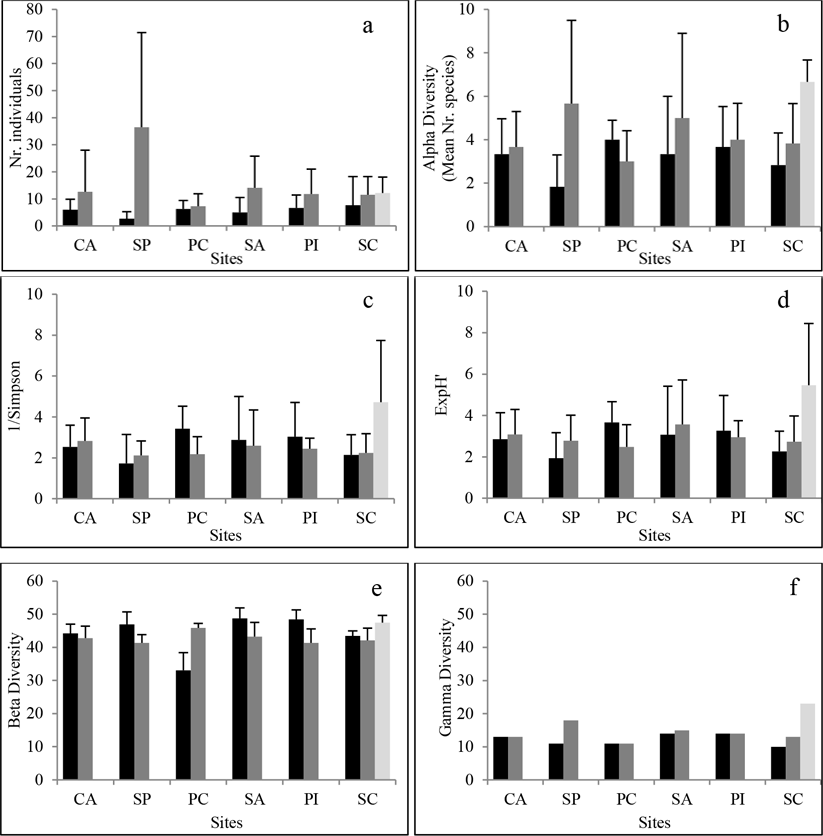
Mean values ± SD of diversity index. a: number of individuals, b: alpha diversity (mean number of species); c: reciprocal Simpson; d: Exponential Shannon; e: beta diversity (% of unshared species); f: gamma diversity (total number of species).

*Cystoseira crinita* at Scannella showed the highest alpha diversity (mean ± SD) of 6 species ± 3, *Cystoseira amentacea* showed the highest alpha diversity at SP (7 ± 4), finally at PC *C. compressa* reached the highest alpha diversity (4 ± 1). The lowest alpha diversity was associated respectively to *C. compressa* at SP with 2 species ± 1 and *C. amentacea* at PC (3 ± 1). The reciprocal Simpson’s as well as the Exponential Shannon showed significant differences between algae but not between sites (Table 4). The data suggested highest diversity in terms of ‘effective number of species’ at SC for *C. crinita* (ExpH’ = 6 ± 3, 1/Simpson = 5 ± 3), the lowest diversity was found at SP for *C. compressa* (ExpH’ = 2 ±, 1/Simpson = 2 ±1). Beta diversity did not show a significant difference among algae and sites (PERMDISP F_12,65_ = 1.5, p > 0.05). Gamma diversity or species richness within a site ranged from 23 species at SC associated with *C. crinita,* to 18 species at SP associated with *C. amentacea* and 14 species at SA and PI associated with *C. compressa,* the lowest gamma diversity corresponded to *C. compressa* at SC with 10 species (Figure 3 a-f).

Two different types of k-dominance curves were plotted, considering or not *M. galloprovincialis Figure* 4.

**Figure 4.**
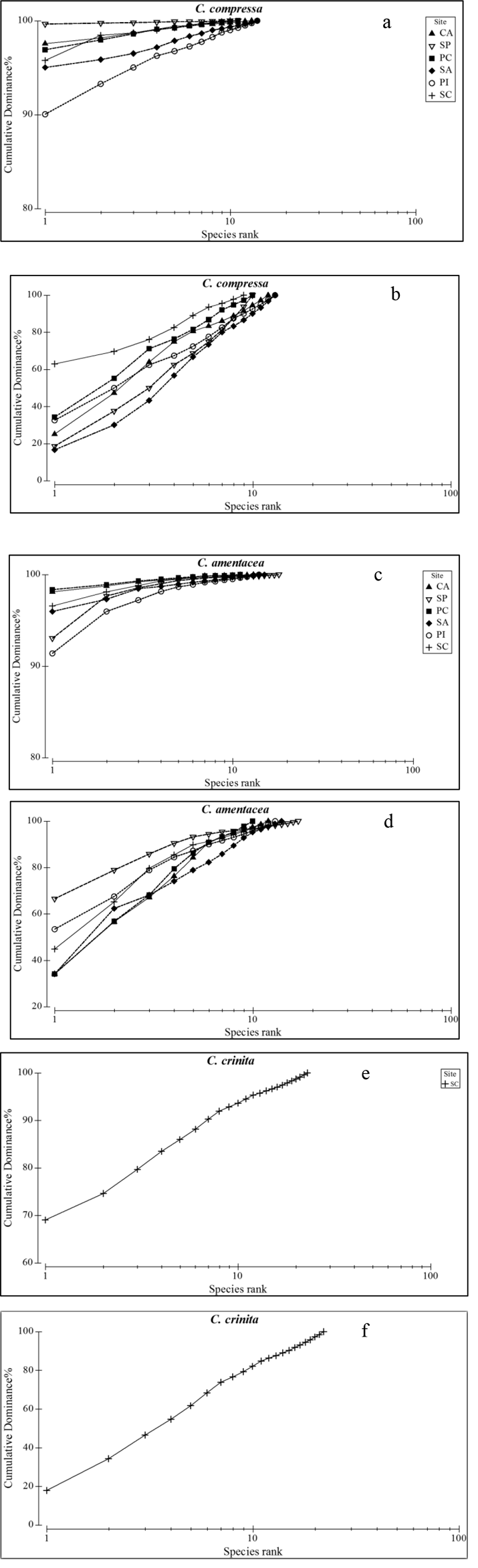
k-dominance curves considering (a, c, e) or not (b, d, f) *M. galloprovincialis.*

It was clear that the species *M. galloprovincialis* dominated over the others in the three algal assemblages at the six sites with high initial value of dominance and k-dominance curves reaching quickly the asymptote.

Excluding *M. galloprovincialis*, in general the three algae hosted diversified molluscs assemblages in the six sites with low initial dominance and k-dominance curves reaching slowly the asymptote, except for *C. compressa* at SC and *Cystoseira amentacea* at SP where the species *Scissurella costata* (63% of total abundance) and *Doto rosea* (67% of total abundance) were the dominant ones respectively. Overall the mollusc community structure differed significantly both among algae and sites (Figure 5). There were significant differences both among algae (F_2,65_ = 7.27, p < 0.01) and sites (F_5,65_ = 3.28, p < 0.001) with a maximum distance between *C. compressa* / *C. crinita.*

**Figure 5.**
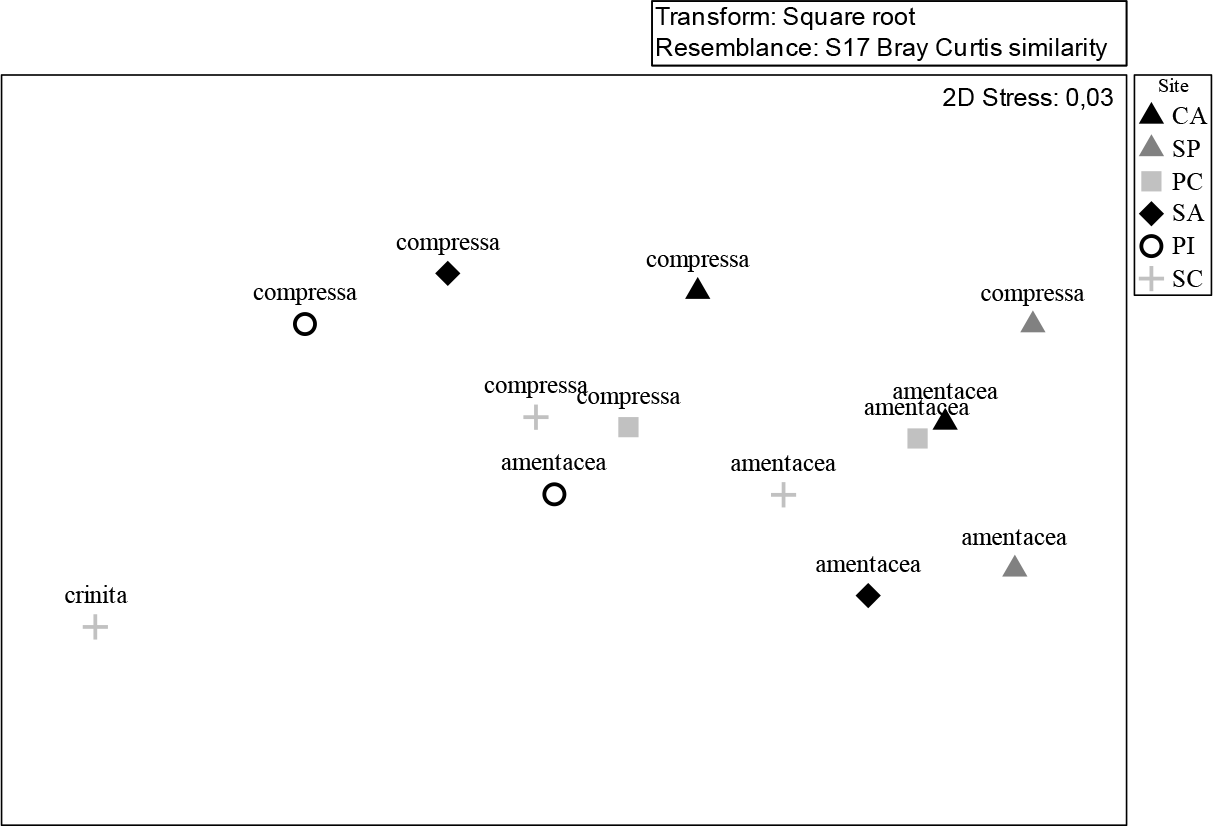
Non-metric multidimensional scaling (nMDS) plot showing the spatial pattern of similarity of mollusc assemblages associated with the three algal species at the six sites performed on the distance among centroids matrix derived from a Bray-Curtis similarity matrix using square-root-transformed abundance data.

The SIMPER analyses highlighted that *C. amentacea* and *C. crinita* displayed an higher average similarity in species composition (28% and 27% respectively) respect to *C. compressa* (17%). The number of species contributing to the 90% of the similarity between the three algal assemblages are 6 for *C. compressa* and *C. crinita* and 5 for *C. amentacea. Scissurella costata* was the most important species in term of percentage of similarity within *C. compressa* and *C. crinita* (46% and 38% respectively), *Doto rosea* contribute with the 54% of similarity for what concern *C. amentacea.* The highest dissimilarity was found among *C. compressa* and *C. crinita* (Table 7)

**Table 7.**
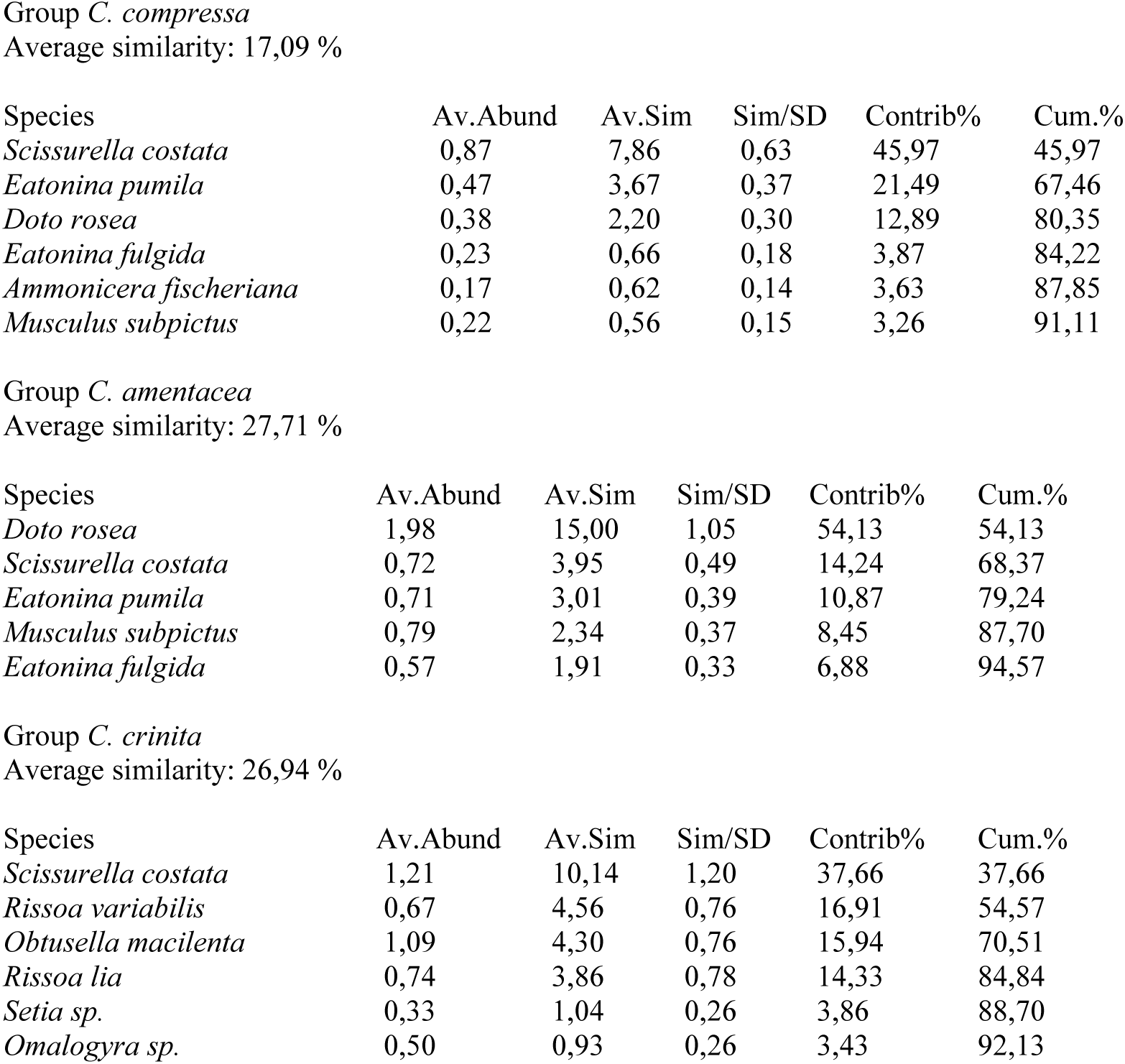
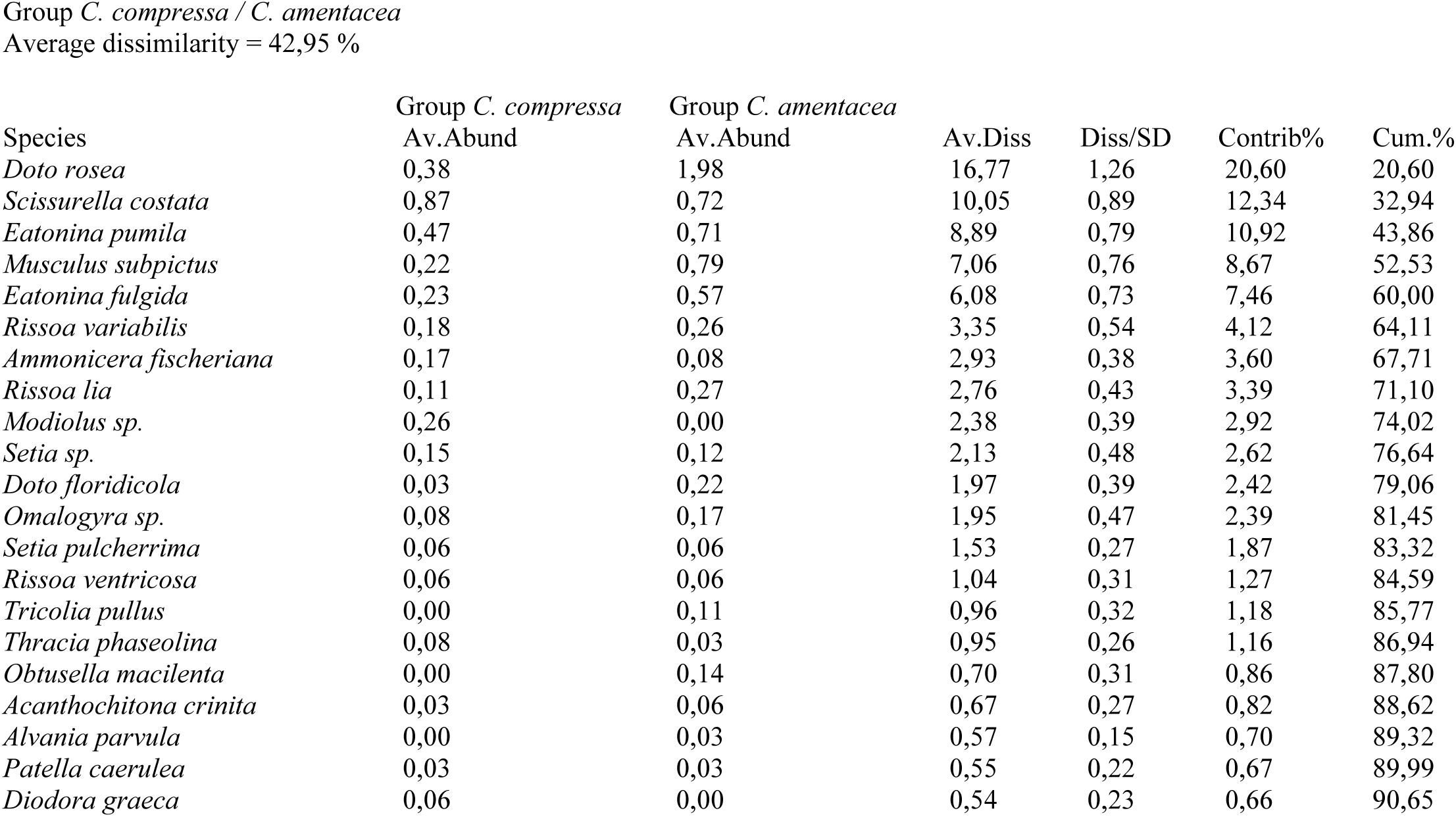
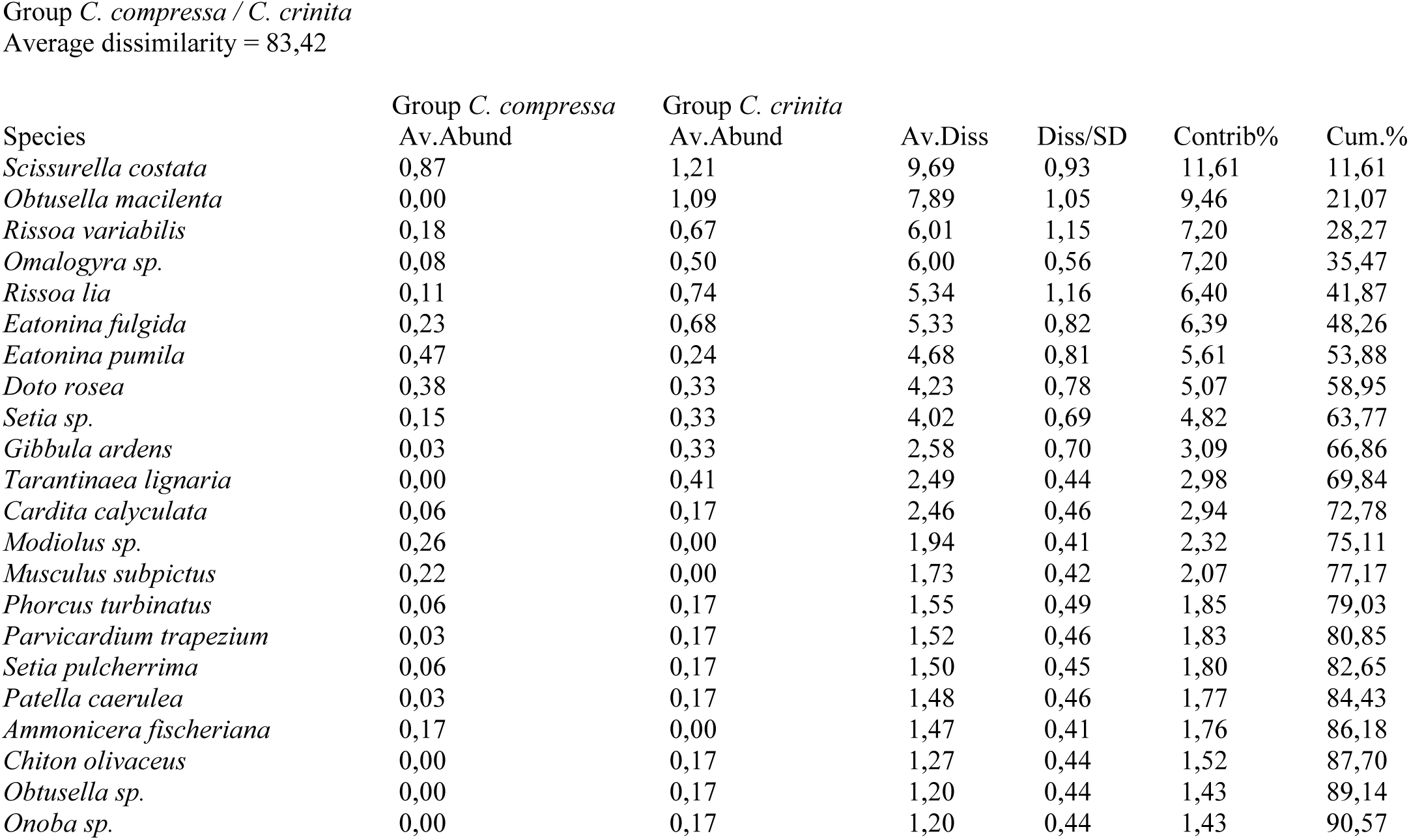
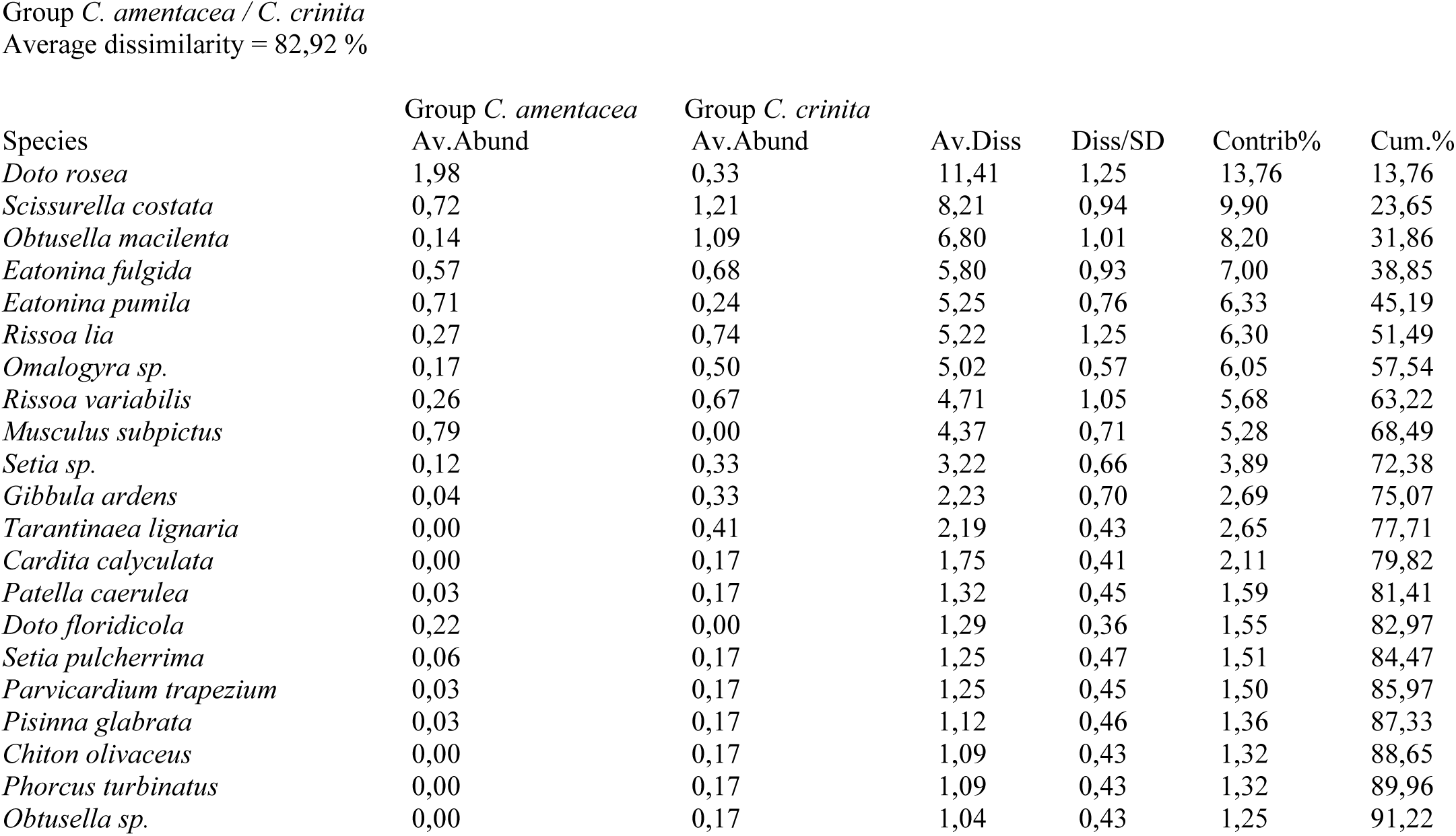
Similarity percentage (SIMPER) results

## Discussion

Macrophytes are important primary producers along coasts worldwide serving as habitat or functioning as ecological engineering species. The beds of seagrasses, kelp and fucoids support epiphytic algae and animals, as well as a variety of associated vagile fauna (Christie et al. 2009). Macroalgae of the genus *Cystoseira* are important engineering species along the temperate rocky coasts all over the Mediterranean sea. Their architectural complex tridimensional structure serve as habitat for a wide variety of organisms both vertebrates and invertebrates (Chemello and Milazzo 2002; Cheminée et al. 2013; Gozler et al. 2010; Pitacco et al. 2014; Urra et al. 2013).

The disappearance of *Cystoseira* species is a phenomenon that has been described in different areas of the Mediterranean Sea, also in the Gulf of Naples (Buia et al. 2013; Grech et al. 2015; Mangialajo et al. 2008; Thibaut et al. 2015; Thibaut et al. 2005). The decline of these habitat forming species also implies the loss of the associated faunal biodiversity. Studies regarding the diversity of invertebrate fauna associated with the seagrass meadow *Posidonia oceanica* at Ischia Island have been the aim of several studies (Gambi 2002; Gambi et al. 1992; Garrard 2013; Mazzella et al. 1992; Mazzella et al. 1989; Scipione et al. 1996; Vasapollo 2009). Our study aims to fill the gap of knowledge concerning the invertebrate fauna composition and diversity associated with the engineering macroalgae of the genus *Cystoseira* along the coasts of Ischia Island where belts of these algae are still present and since previously have never been assessed. We have analyzed the molluscs community associated with the three *Cystoseira* assemblages, *Cystoseira compressa, Cystoseira amentacea* and *Cystoseira crinita* along the coasts of Ischia Island and the pattern of diversity at a small geographic spatial scale.

It is known that different macroalgae do not support benthic fauna in the same way (Williams and Seed 1992) and this may depend on several factors such as the life cycles, algal architecture or the exhibition of chemical defenses (Duffy and Hay 1994). Different algal shapes are important in determining patterns of abundance and size structure of the associated fauna (Edgar 1983).

The overall pattern of spatial distribution of these three algal species and their main architectural attributes are quite different within the sampling sites taken into account for the present study. *Cystoseira compressa* was the more widespread species at the six sampling site holding dense and continuous belts, *Cystoseira amentacea* assemblages were dense but more scattered over the sites, *Cystoseira crinita* only occurred in a tide pool close to the sea surface at Scannella where it completely covers the rocky pool walls. Although *Cystoseira compressa* reached the highest mean values of density in all the sites compared to *Cystoseira amentacea*, it is characterized by shorter and less branched thalli. The biomass does not show significant differences among the three algal assemblages. The height represents thus the most important macroalgal feature in diversifying the algal associations among the six sampling sites.

Although the number of mollusc species associated with *Cystoseira amentacea* and *Cystoseira compressa* was the same, *C. amentacea* hosts an higher number of individuals at the six sampling sites, probably because the longer thalli of the latter algal species offers a wider surface of colonization. The maximum total number of species (gamma diversity) at local scale as well as the maximum mean number of species per sampling unit (alpha diversity) was found at Scannella for the species *C. crinita*, this data was confirmed by the highest values of the diversity index too. The number of individuals associated with *C. crinita* at SC was comparable to that of the other two algal species at the same site, this data together with the high value of beta diversity (percentage of unshared species within the sampling site) seems to highlight that *C. crinita* malacofauna was more heterogeneous in terms of species composition (a higher number of different species most of which are unshared among the different sampling units). At Scannella the species *C. crinita* has longer thalli than those of *C. compressa* and *C. amentacea,* this could be related to the peculiar habitat of the rocky pool representing a sheltered zone with low hydrodynamic regime and a low competition for space with the other algal species. Apart from *C. crinita* at SC, the highest values of alpha diversity are due to *C. amentacea* at all the sampling sites although the beta diversity is lower in *C. amentacea* respect to *C. compressa.* This data seems to suggest that *C. amentacea* malacofauna is more homogeneous in terms of species composition among the different sites.

A total of 53 mollusc species were identified hosted by the three *Cystoseira* species along the coast of Ischia Island. Gastropoda represents the dominant taxa in terms of number of species followed by Bivalvia and Polyplacophora. This trend is confirmed by other studies on the molluscs assemblage associated with photophilous algal stands in other areas of the Mediterranean Sea (Antoniadou and Chintiroglou 2005; Chemello and Milazzo 2002; Pitacco et al. 2014; Poulicek 1985; Sánchez-Moyano et al. 2000). The most species-rich family was Rissoidae, two species are exclusively associated with *C. amentacea* as well as two other species are exclusively associated with *Cystoseira crinita,* no species of Rissoidae are exclusively associated with *Cystoseira compressa*. These species are micrograzers feeding preferentially on diatoms and epiphyte microalgae laying on *Cystoseira* leaves. Some other species are exclusively associated with only one or two algal associations as shown in the Table 4.

The most frequent and top dominant mollusc species inhabiting *Cystoseira* associations along the coast of Ischia Island was the bivalve *Mytilus galloprovincialis* (96.6 % of the total abundance of all the individuals). However only the juvenile stages (the most represented size ranges among the 0.3 – 3 mm) were strictly associated with algal canopy. The adult individuals were mostly found under the algal canopies attached to the hard rocky bottoms or among the holdfasts of macroalgae where they construct a real continuous barrier competing for the space with the above algal associations. Except the presence of the mussels, it is possible to identify some differences in the pattern of association of molluscs community within the three algal assemblages, although the low level of dominance. In general, the three algal species hosted diversified molluscs assemblages at the six sampling sites (Figure 5) with low initial dominance and k-dominance curves reaching slowly the asymptote (Figure 4 A-F). The gastropod *Scrissurella costata* is the most frequent and abundant species associated with *Cystoseira compressa*; this species is a deposit feeder feeding on food trapped in sediments retained by algal thalli. This association could be related to the high density of this algal species at all the analyzed sampling sites where it creates a tangled layer with its holdfast trapping high quantity of sediments. *Scissurella costata* was also the most frequent species associated with *Cystoseira crinita* at Scannella, this alga-animal association could be related to the peculiar habitat of rocky pool in which these species thrive. *Cystoseira crinita* in fact usually grows in the upper infralittoral zone on both low and intermediately exposed gently sloping rocky bottoms that are often subjected to a high degree of sedimentation (Sales and Ballesteros 2009; Sales and Ballesteros 2010). *Doto rosea* was the most frequent and abundant species associated with *Cystoseira amentacea*, this species is a specialized carnivore that commonly lives on hydroids, probably this species is more associated to the epibiontic hydroids living on *Cystoseira* surface rather than directly to the alga.

The highest dissimilarity in terms of molluscs composition was found among *C. compressa* and *C. crinita*, while *C. compressa* and *C. amentacea* are more similar.

The analyzed three algal associations only host juvenile molluscs stages, no adults were found. This confirms the importance of *Cystoseira* species associations as a nursery for molluscs recruitment.

No significant differences were found in the composition and the number of species at the different sampling sites, this let to suppose that the occurrence of the different zones within the marine protected area Regno di Nettuno does not influence the pattern of molluscs biodiversity associated with these algal species.

Comparing results from different areas is a challenging task because of the natural variability among geographic zones and the different sampling methods used. Although these difficulties, some similarities could be found in the number of mollusc species associated with upper infralittoral *Cystoseira* associations from other sites of the Mediterranean Sea. For example Chemello and Milazzo (2002) reported 35 species of molluscs associated with the species *Cystoseira barbatula* and *Cystoseira spinosa* at a shallow rocky shore at Lampedusa Island. Pitacco et al. (2014) reported 69 species of molluscs associated with the two algal sub-associations of *Cystoseiretum crinitae* and the association *Cystosereitum barbate* at the Gulf of Trieste. Çulha et al. (2010) found a total of 14 species associated with *Cystoseira barbata* faces at the Sinop Peninsula (southern Black Sea). Gozler et al. (2010) recorded 7 molluscs species associated with the species *Cystoseira barbata* at the southeastern Black Sea.

Although the dominance of the bivalve *Mytilus galloprovincialis* at all the analyzed sampling sites, the three species of *Cystoseira* are able to support diversified and structured molluscs assemblages. These results confirm the importance of *Cystoseira* associations in structuring habitat eligible for the mollusc assemblages especially during the juvenile stages. These results must be taken as an incentive for a series of protection strategies towards these important habitat forming species since these are able to serve as a nursery and sheltered habitat supporting therefore a good level of associated biodiversity.

## Acknowledgments

This work has been partially funded by the Italian Flagship Project RITMARE – The Italian Research for the Sea. RITMARE also supported A. Chiarore’s grant. We thank V. Rando and B. Iacono for their support in the fieldwork.

